# Flavin Monooxygenases Regulate *C. elegans* Axon Guidance and Growth Cone Protrusion with UNC-6/Netrin signaling and Rac GTPases

**DOI:** 10.1101/131755

**Authors:** Mahekta Gujar, Aubrie M. Stricker, Erik A. Lundquist

## Abstract

The guidance cue UNC-6/Netrin regulates both attractive and repulsive axon guidance. Our previous work showed that in *C. elegans*, the attractive UNC-6/Netrin receptor UNC-40/DCC stimulates growth cone protrusion, and that the repulsive receptor, an UNC-5/UNC-40 heterodimer, inhibits growth cone protrusion. We have also shown that inhibition of growth cone protrusion downstream of the UNC-5/UNC-40 repulsive receptor involves Rac GTPases, the Rac GTP exchange factor UNC-73/Trio, and the cytoskeletal regulator UNC-33/CRMP, which mediates Semaphorin-induced growth cone collapse in other systems. The multidomain flavoprotein monooxygenase (FMO) MICAL also mediates growth cone collapse in response to Semaphorin by directly oxidizing F-actin, resulting in depolymerization. The *C. elegans* genome does not encode a multidomain MICAL-like molecule, but does encode five flavin monooxygenases (FMO-1, -2, -3, -4, and 5) and another molecule, EHBP-1, similar to the non-FMO portion of MICAL.

Here we show that FMO-1, FMO-4, FMO-5, and EHBP-1 may play a role in UNC-6/Netrin directed repulsive guidance mediated through UNC-40 and UNC-5 receptors. Mutations in *fmo-1, fmo-4, fmo-5,* and *ehbp-1* showed VD/DD axon guidance and branching defects, and variably enhanced *unc-40* and *unc-5* VD/DD guidance defects. Developing growth cones *in vivo* of *fmo-1, fmo-4, fmo-5,* and *ehbp-1* mutants displayed excessive filopodial protrusion, and transgenic expression of FMO-5 inhibited growth cone protrusion. Mutations suppressed growth cone inhibition caused by activated UNC-40 and UNC-5 signaling, and activated Rac GTPase CED-10 and MIG-2, suggesting that these molecules are required downstream of UNC-6/Netrin receptors and Rac GTPases. From these studies, we conclude that FMO-1, FMO-4, FMO-5, and EHBP-1 represent new players downstream of UNC-6/Netrin receptors and Rac GTPases that inhibit growth cone filopodial protrusion in repulsive axon guidance.

**Author Summary:** Molecular mechanisms of axon repulsion mediated by UNC-6/Netrin are not well understood. Inhibition of growth cone lamellipodial and filopodial protrusion is critical to repulsive axon guidance. Previous work identified a novel pathway involving Rac GTPases and the cytoskeletal interacting molecule UNC-33/CRMP required for UNC-6/Netrin-mediated inhibition of growth cone protrusion. In other systems, CRMP mediates growth cone collapse in response to semaphorin. Here we demonstrate a novel role of flavoprotein monooxygenases (FMOs) in repulsive axon guidance and inhibition of growth cone protrusion downstream of UNC-6/Netrin signaling and Rac GTPases. In *Drosophila* and vertebrates, the multidomain MICAL FMO mediates semaphorin-dependent growth cone collapse by direct oxidation and depolymerization of F-actin. The *C. elegans* genome does not encode a multidomain MICAL-like molecule, and we speculate that the *C. elegans* FMOs might have an equivalent role downstream of UNC-6/Netrin signaling. Indeed, we show that EHBP-1, similar to the non-FMO portion of MICAL, also controls repulsive axon guidance and growth cone inhibition, suggesting that in *C. elegans*, the functions of the multidomain MICAL molecule might be distributed across different molecules. In sum, we show conservation of function of molecules involved in semaphorin growth cone collapse with inhibition of growth cone protrusion downstream of UNC-6/Netrin signaling.

## Introduction

The formation of neural circuits during development depends on the guidance of growing axons to their proper synaptic targets. This process relies on the growth cone, a dynamic actin based structure present at the tip of a growing axon. Growth cones contain a dynamic lamellipodial body ringed by filopodial protrusions, both important in guiding the axon to its target destination [1-4]. Guidance receptors present on the leading edge of the growth cone sense and respond to various extracellular guidance cues, which attract or repel axons enabling them to reach their proper target destination [5, 6].

The secreted laminin-like guidance molecule UNC-6/Netrin mediates both axon attraction and axon repulsion and defines a dorsal-ventral guidance mechanism conserved from invertebrates to vertebrates [7-9]. Attractive or repulsive responses to UNC-6/Netrin depend on the receptors expressed on the growth cone. Homodimers of the UNC-6/Netrin receptor UNC-40/DCC mediate attraction, and UNC-5-UNC-40 heterodimers or UNC-5 homodimers mediate repulsion [10-12].

In *C. elegans,* UNC-6/Netrin is secreted by the ventral cells and along with its receptors UNC-40 and UNC-5 is required for the dorsal ventral guidance of circumferential neurons and axons [8, 13, 14]. Previous studies of repelled VD growth cones in Netrin signaling mutants revealed a correlation between attractive axon guidance and stimulation of growth cone protrusion, and repulsive axon guidance and inhibition of growth cone protrusion [15]. For example, in *unc-5* mutants, growth cones were larger and more protrusive, and often displayed little or no directed movement. This is consistent with observation that increased growth cone size was associated with decreased neurite growth length [16]. Conversely, constitutive activation of UNC-40/UNC-5 signaling in repelled VD growth cones led to smaller growth cones with severely reduced filopodial protrusion [15, 17]. Thus, directed growth cone repulsion away from UNC-6/Netrin requires a balance of pro- and anti-protrusive activities of the receptors UNC-40 and UNC-40-UNC-5, respectively, in the same growth cone [15].

Genetic analysis has identified a cytoskeletal signaling pathway involved in stimulation of growth cone protrusion in response to the attractive UNC-40 signaling that includes CDC-42, the Rac-specific guanine nucleotide exchange factor TIAM-1, the Rac-like GTPases CED-10 and MIG-2, as well as the cytoskeletal regulators Arp2/3 and activators WAVE-1 and WASP-1, UNC-34/Enabled, and UNC-115/abLIM [18-23], consistent with findings in other systems [7]. Mechanisms downstream of UNC-5 in axon repulsion are less well described, but the PH/MyTH4/FERM molecule MAX-1 and the SRC-1 tyrosine kinase have been implicated [24, 25]. We delineated a new pathway downstream of UNC-5 required for its inhibitory effects on growth cone protrusion, involving the Rac GEF UNC-73/Trio, the Rac GTPases CED-10 and MIG-2, and the cytoskeletal-interacting molecule UNC-33/CRMP [17].

Collapsin response mediating proteins (CRMPs) were first identified as mediators of growth cone collapse in response to the Semaphorin family of guidance cues [26], and we have shown that UNC-33/CRMP inhibits growth cone protrusion in response to Netrin signaling [17]. This motivated us to consider other mediators of Semaphorin-induced growth cone collapse in Netrin signaling. In *Drosophila*, the large multidomain cytosolic protein MICAL (Molecule Interacting with CasL) is required for the repulsive motor axon guidance mediated by interaction of Semaphorin 1a and Plexin A [27, 28]. MICAL proteins are a class of flavoprotein monooxygenase enzymes that bind flavin adenine dinucleotide (FAD) and use the cofactor nicotinamide dinucleotide phosphate (NADPH) to facilitate oxidation-reduction (Redox) reactions [27]. MICAL regulates actin disassembly and growth cone collapse in response to semaphorin via direct redox interaction with F-actin [29, 30]. MICAL molecules from *Drosophila* to vertebrates have a conserved domain organization: and N-terminal flavin-adenine dinucleotide (FAD)-binding monooxygenase domain, followed by a calponin homology (CH) domain, a LIM domain, a proline-rich domain, and a coiled-coil ERM α-like motif [27, 31].

The *C. elegans* genome does not encode for a MICAL-like molecule with the conserved domain organization described above. However, it does contain five flavin monooxygenase (*fmo*) genes similar to the Flavin monooxygenase domain of MICAL: *fmo-1, fmo-2, fmo-3, fmo-4 and fmo-5* [32]. Like MICAL, the *C. elegans* FMO molecules contain an N-terminal FAD binding domain and a C-terminal NADP or NADPH binding domain [27, 32]. The *C. elegans* gene most similar to the non-FMO portion of MICAL is the Eps-15 homology domain binding protein EHBP-1 [33], which contains a CH domain as does MICAL.

In this work, we test the roles of the *C. elegans* FMOs and EHBP-1 in Netrin-mediated axon guidance and growth cone protrusion. We find that *fmo-1, fmo-4, fmo-5* and *ehbp-1* mutants display pathfinding defects of the dorsally-directed VD/DD motor neuron axons that are repelled by UNC-6/Netrin, and that they interact genetically with *unc-40* and *unc-5*. We also find that VD growth cones in these mutants display increased filopodial protrusion, similar to mutants in repulsive UNC-6/Netrin signaling (e.g. *unc-5* mutants), and that transgenic expression of FMO-5 inhibits growth cone protrusion, similar to constitutively-activated UNC-40 and UNC-5. We also show that FMO-1, FMO-4, FMO-5 and EHBP-1 are required for the growth cone inhibitory effects of activated UNC-5, UNC-40, and the Rac GTPases CED-10 and MIG-2. Together, these genetic analyses suggest that FMO-1, FMO-4, FMO-5, and EHBP-1 normally restrict growth cone protrusion, and that they might do so in UNC-6/Netrin-mediated growth cone repulsion.

## Results

### *fmo-1,4,5* and *ehbp-1* affect VD/DD axon pathfinding

The *C. elegans* genome lacks an apparent homolog of MICAL. However, it contains five flavin monooxygenase genes (*fmo-1,2,3,4,*5) (Figure 1A) [32]. The *C. elegans* molecule most similar to the non-FMO portion of MICAL is EHBP-1, the homolog of the mammalian EH domain binding protein 1 (Ehbp1) protein [33]. We analyzed existing mutations in *fmo* genes and *ehbp-1* (Figure 1B) for VD/DD axon guidance defects*. fmo-1(ok405)* was a 1,301-bp deletion that removed part of exon 3 and all of exons 4, 5 and 6. *fmo-2(ok2147)* was a 1070-bp deletion that removed part of exon 4 and 5.*fmo-4(ok294)* was a 1490-bp deletion that removed all of exons 2, 3, 4 and 5. *fmo-5(tm2438)* is a 296-bp deletion which removes part of intron 3 and exon 4. These deletions all affected one or both predicted enzymatic domains of the FMO molecules. *fmo-3(gk184651)* was a G to A substitution in the 3’ splice site of intron 6. *ehbp-1(ok2140)* is a 1,369-bp deletion that removed all of exon 5 and 6.

**Fig. 1.**
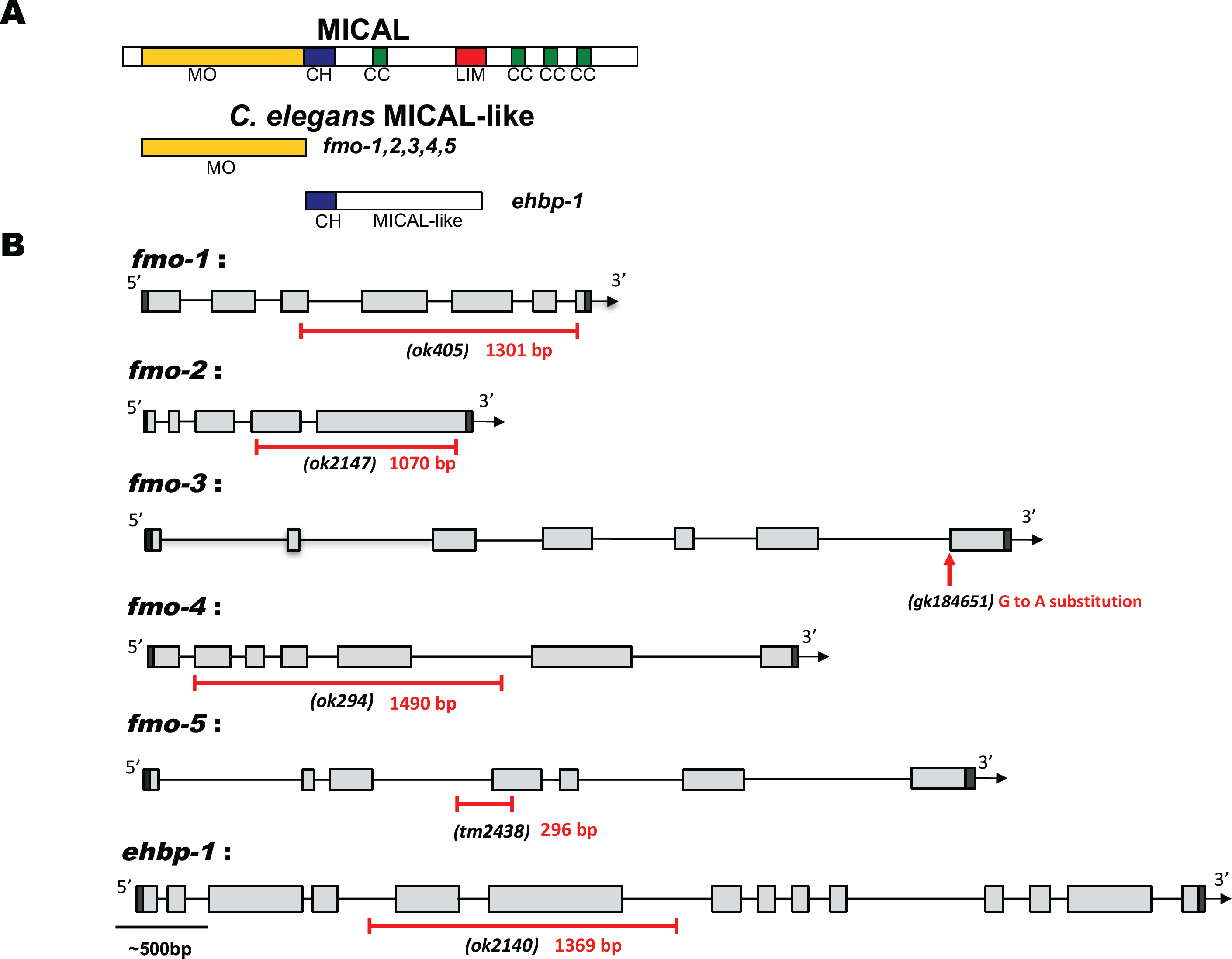
*fmo* genes and *ehbp-1*. **(A)** Diagram of Drosophila MICAL, the *C. elegans* flavin monooxygenases (FMOs), and *C. elegans* EHBP-1. MO, flavin monooxygenase domain; CH-calponin homology domain, CC, coiled coil domain; LIM, LIM domain. **(B)** The structures of the *fmo-1, fmo-2, fmo-3, fmo-4, fmo-5* and *ehbp-1* genes are shown. Filled boxes represent exons. The extent of deletions in *fmo-1, fmo-2, fmo-4, fmo-5* and *ehbp-1* are shown below the structure, indicated by a red line. The red arrow points to the region of the splice site mutation in *fmo-3*. Scale bar indicates 500bp.

The 19 D-class motor neurons cell bodies reside in the ventral nerve cord. They extend axons anteriorly and then dorsally to form a commissure, which normally extend straight dorsally to the dorsal nerve cord (Figure 2 and Figure 3B) On the right side of wild-type animals, an average of 16 commissures were observed, due to the fasciculation of some processes as a single commissure (Figure 2C and Materials and Methods). *fmo-1,4* and *5* and *ehbp-1* mutants showed significant defects in VD/DD axon pathfinding, including ectopic axon branching and wandering (∼3-5%; see Materials and Methods and Figure 3A, C and D). *fmo-2* and *fmo-3* mutations showed no significant defects compared to *wild-type* (Figure 3A). Most double mutants showed no strong synergistic defects compared to the predicted additive effects of the single mutants (Figure 3E). However, the *fmo-2; fmo-3* and the *fmo-2; fmo-4* double mutants showed significantly more defects compared to the predicted additive effects of the single mutants. The *fmo-4; ehbp-1* double mutant displayed significantly reduced defects than either mutation alone. Lack of extensive phenotypic synergy suggests that the FMOs do not act redundantly, but rather that they might have discrete and complex roles in axon guidance, as evidenced by *fmo-4; ehbp-1* mutual suppression.

**Fig. 2.**
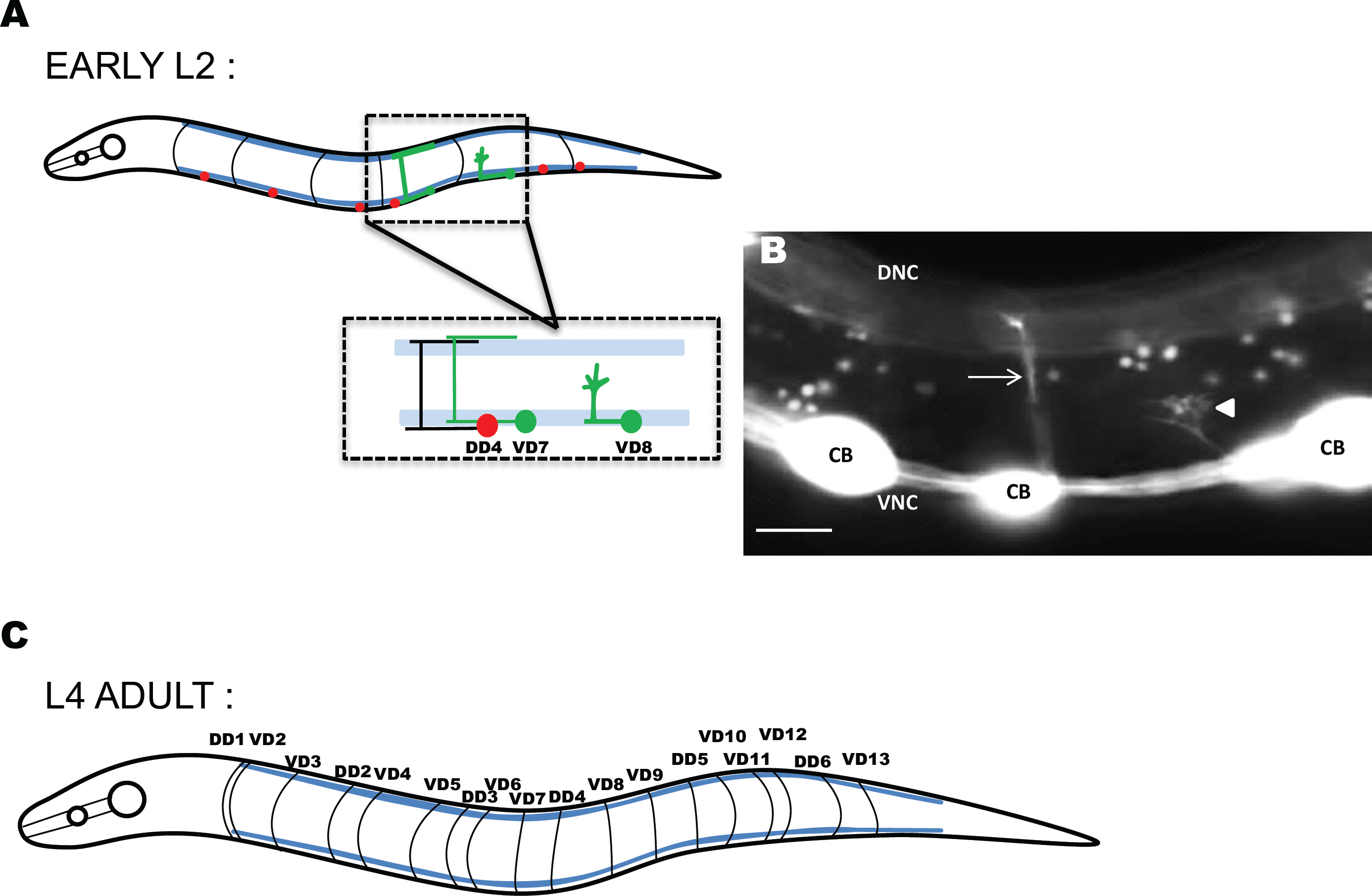
VD/DD motor neurons and axons in *C. elegans*. **(A)** Diagram of an early L2 larval *C. elegans* hermaphrodite highlighting the position and structure of the DD motor neurons (red) and axons (black). Anterior is to the left, and dorsal is up. The blue lines represent the ventral and dorsal muscle quadrants. In the early L2 larval stage, the VD neurons (green) extend axons anteriorly in the ventral nerve cord after which the axons turn dorsally and migrate to the dorsal nerve cord to form commissures. Only two of the 13 VD neurons are shown. While migrating towards the dorsal nerve cord, VD growth cones display an extended, protrusive morphology with highly dynamic filopodial protrusions (VD8). VD7 shows the final structure of the VD neurite. **(B)** Fluorescent micrograph of an early L2 larval wild-type commissure indicated by an arrow, and a VD growth cone indicated by an arrowhead. CB, cell body; DNC, dorsal nerve cord; and VNC, ventral nerve cord. Scale bar represents 5μm. **(C)** Diagram of an L4 hermaphrodite after all the VD axon outgrowth is complete. The 18 commissures on the right side of the animal are shown (black lines), and axon guidance defects of these commissures were scored. One commissure (VD1) extends on the right side and was not scored. Of the 18 commissures on the right side, two (DD1 and VD2) extend as a single fascicle. Others pairs occasionally extended as single fascicles as well, resulting in an average of 16 observable commissures per *wild-type* animal.

**Fig. 3.**
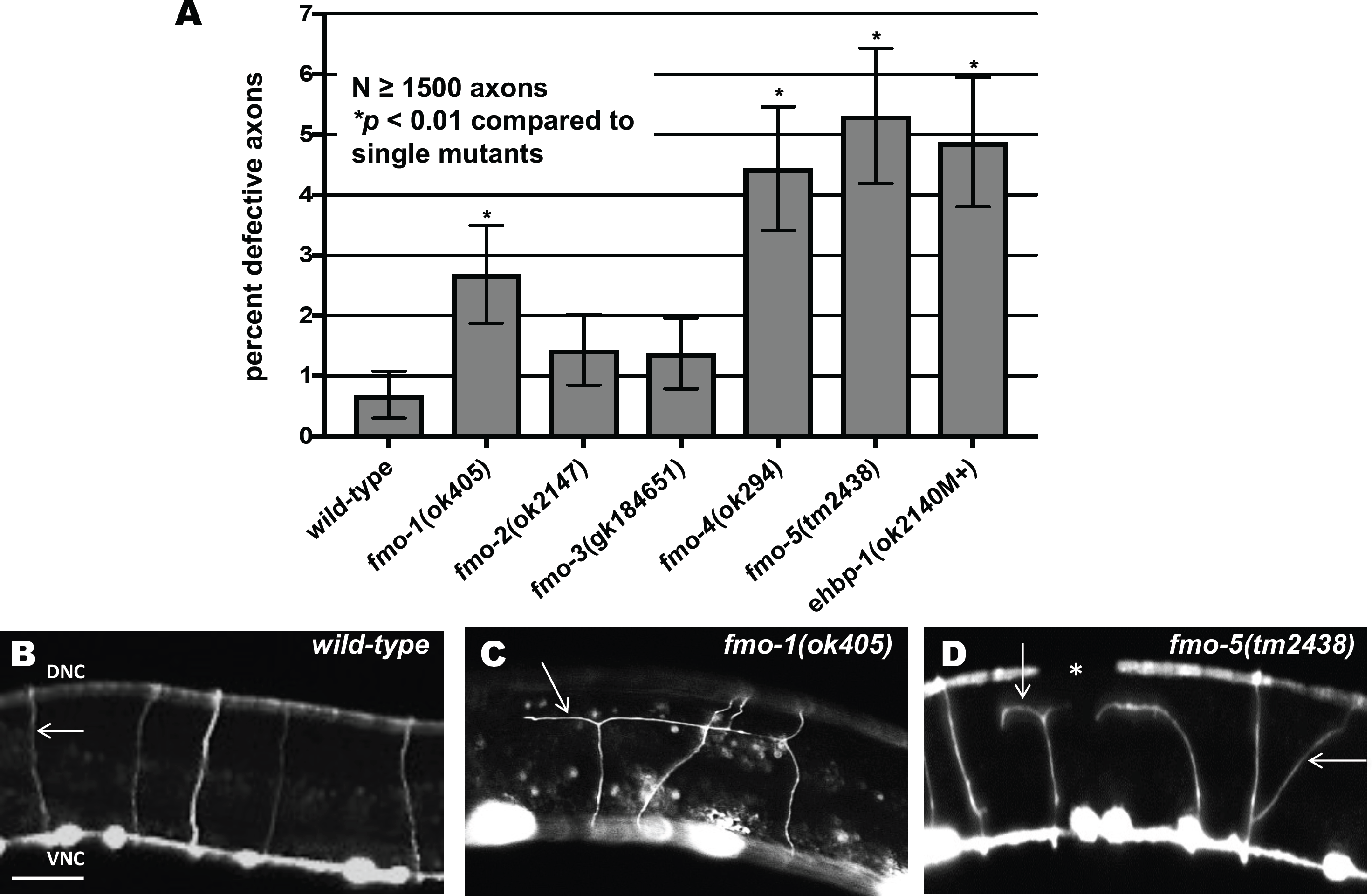
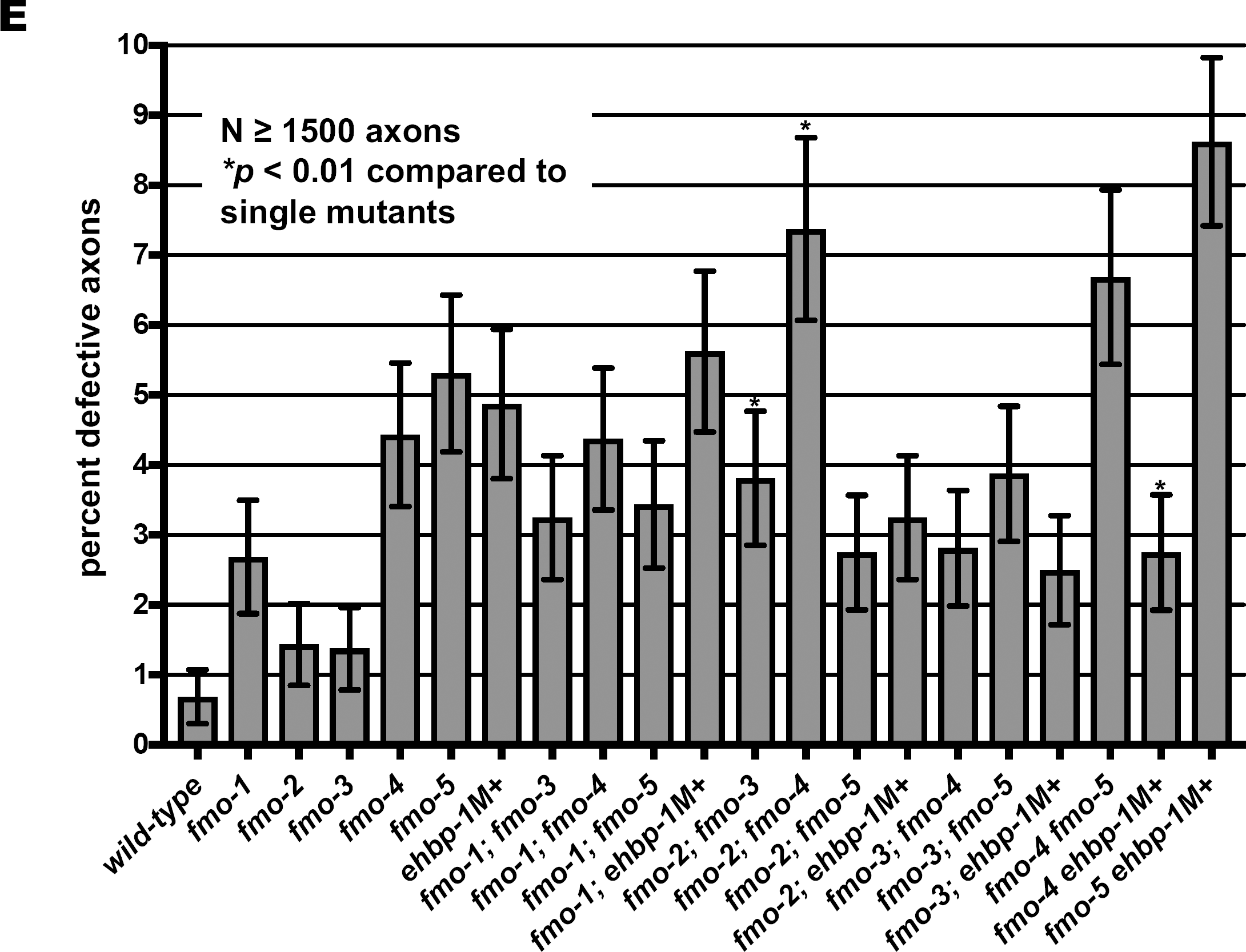
Mutations in *fmo-1, fmo-4, fmo-5* and *ehbp-1* cause axon pathfinding defects. **(A)** Percentage of VD/DD axons with pathfinding defects (see Materials and Methods) in single mutants harboring the *juIs76[Punc-25::gfp]* transgene. Asterisks indicate the significant difference between wild-type and the mutant phenotype (\**p* < 0.01) determined by Fischer’s exact test. Error bars represent 2x standard error of proportion. **(B-D)** Representative fluorescent micrograph of L4 VD/DD axons. Anterior is to the left, and dorsal is up. The scale bar represents 5μm. DNC, dorsal nerve cord; and VNC, ventral nerve cord. (B) A *wild-type* commissure is indicated by an arrow. (C) An *fmo-1(ok405)* commissure branched and failed to reach to dorsal nerve cord (arrow). (D) *fmo-5(tm2438)* VD/DD axons branched and wandered (arrows). A gap in the dorsal nerve cord (asterisk) indicates that commissural processes failed to reach the dorsal nerve cord. **(E)** Percentage of VD/DD axons with pathfinding defects in double mutants as described in (A). Asterisks (*) indicate significant difference between double mutants and the predicted additive effect of single mutants (*p* < 0.01) (see Materials and Methods) determined by Fischer’s exact test. At least 1500 axons were scored per genotype. M+ indicates that the animal has wild-type maternal *ehbp-1(+)* activity.

### Axon pathfinding defects of *unc-40* and *unc-5* are increased by *fmo-1, fmo-4* and *fmo-5* mutations

In *unc-40(n324)* strong loss-of-function mutants, most axons (92%) extended past the lateral midline despite wandering (see Materials and Methods and Figures 4A and B). *fmo-1, fmo-4, fmo-5* and *ehbp-1* displayed < 1% failing to extend past the lateral midline (Figure 4A). *fmo-1, fmo-4,* and *fmo-5* mutations significantly enhanced the VD/DD lateral midline crossing defects of *unc-40(n324)* (Figure 4A and C). *ehbp-1* did not enhance *unc-40* (Figure 4A).

**Fig. 4.**
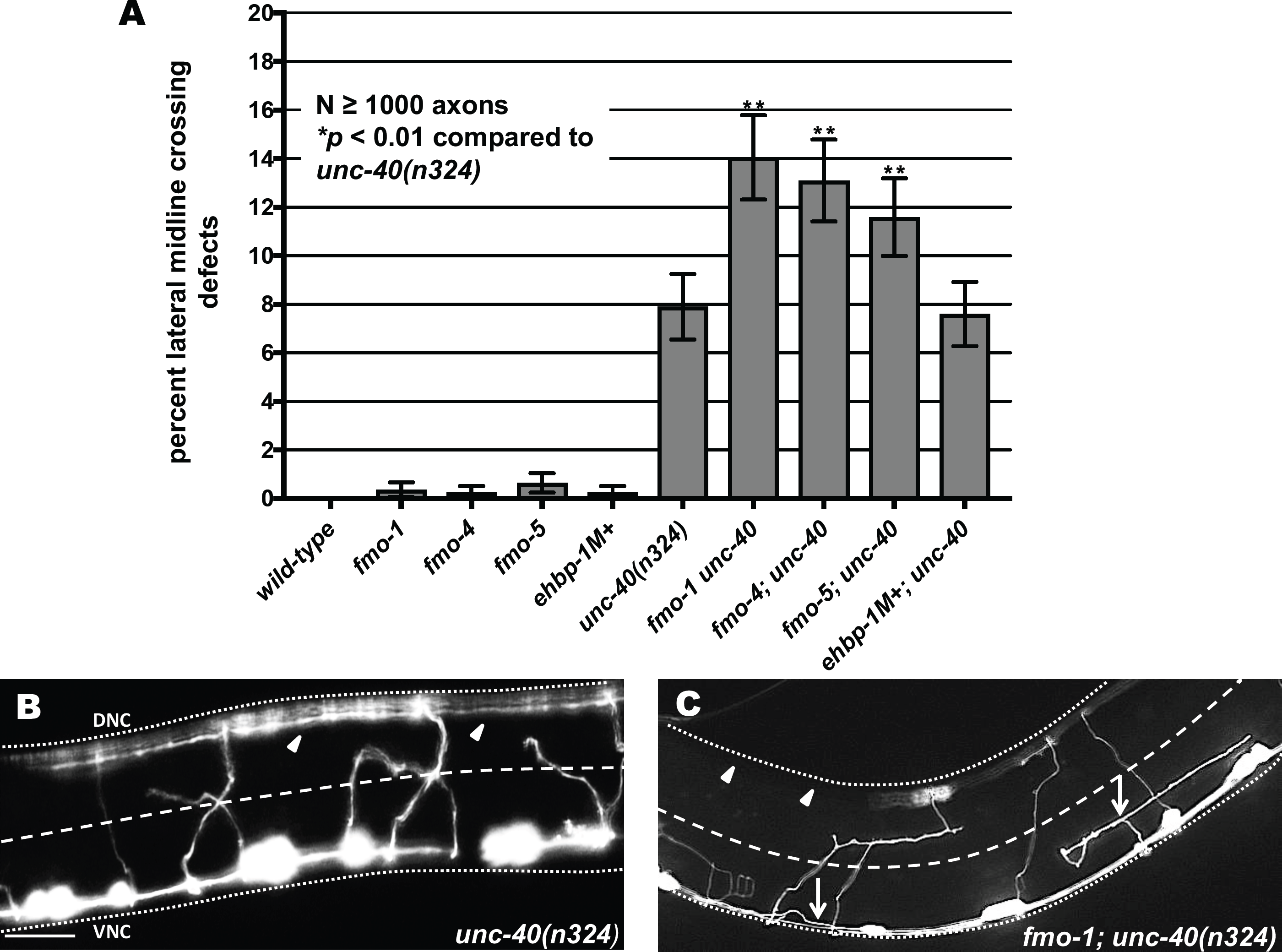
Axon pathfinding defects in *unc-40(n324)* are enhanced by loss of *fmo-1, fmo-4* and *fmo-5*. **(A)** Percentage of VD/DD axons that failed cross the lateral midline of L4 hermaphrodites. Error bars represent 2x standard error of the proportion; double asterisks (**) indicates a significant difference between *unc-40(n324)* alone and the double mutants (*p* < 0.001) determined by Fisher’s exact test. Only axon commissures visibly emanating from the ventral nerve cord were scored. **(B,C)** Representative images showing VD/DD axons (arrows) after their complete outgrowth in L4 animals. The lateral midline of the animal is indicated by the dashed white line. The dorsal nerve cord and ventral nerve cord are indicated by a dotted white line. Dorsal is up, anterior is to the left. Scale bar represents 5μm. (B) In *unc-40(n324)*, many axons extend past the lateral midline, as evidenced by axons in the dorsal nerve cord (arrowheads). (C) In *fmo-1(ok405); unc-40(n324)*, an increased number of axons did not cross the midline resulting in extensive regions of dorsal nerve cord without axons (arrowheads). Arrowhead indicates large gaps in the dorsal nerve cord.

*unc-5(e53)* strong loss-of-function mutants display a nearly complete failure of VD axons to reach the dorsal nerve cord [13, 15]. *unc-5(e152)* is a hypomorphic allele [34] and displayed 22% failure of axons to cross the lateral midline (Figure 5A). The *unc-5(op468)* allele [35] also displayed a weaker lateral midline crossing phenotype (10%), indicating that it is also a hypomorphic allele (Figure 5B). *fmo-1, fmo-4* and *fmo-*5 significantly enhanced the VD/DD axon guidance defects of both *unc-5(e152)* and *unc-5(op468)*, but *ehbp-1* did not (Figure 5). These results indicate that FMO-1,4, and 5 might act with UNC-40 and UNC-5 in VD/DD axon pathfinding.

**Fig. 5.**
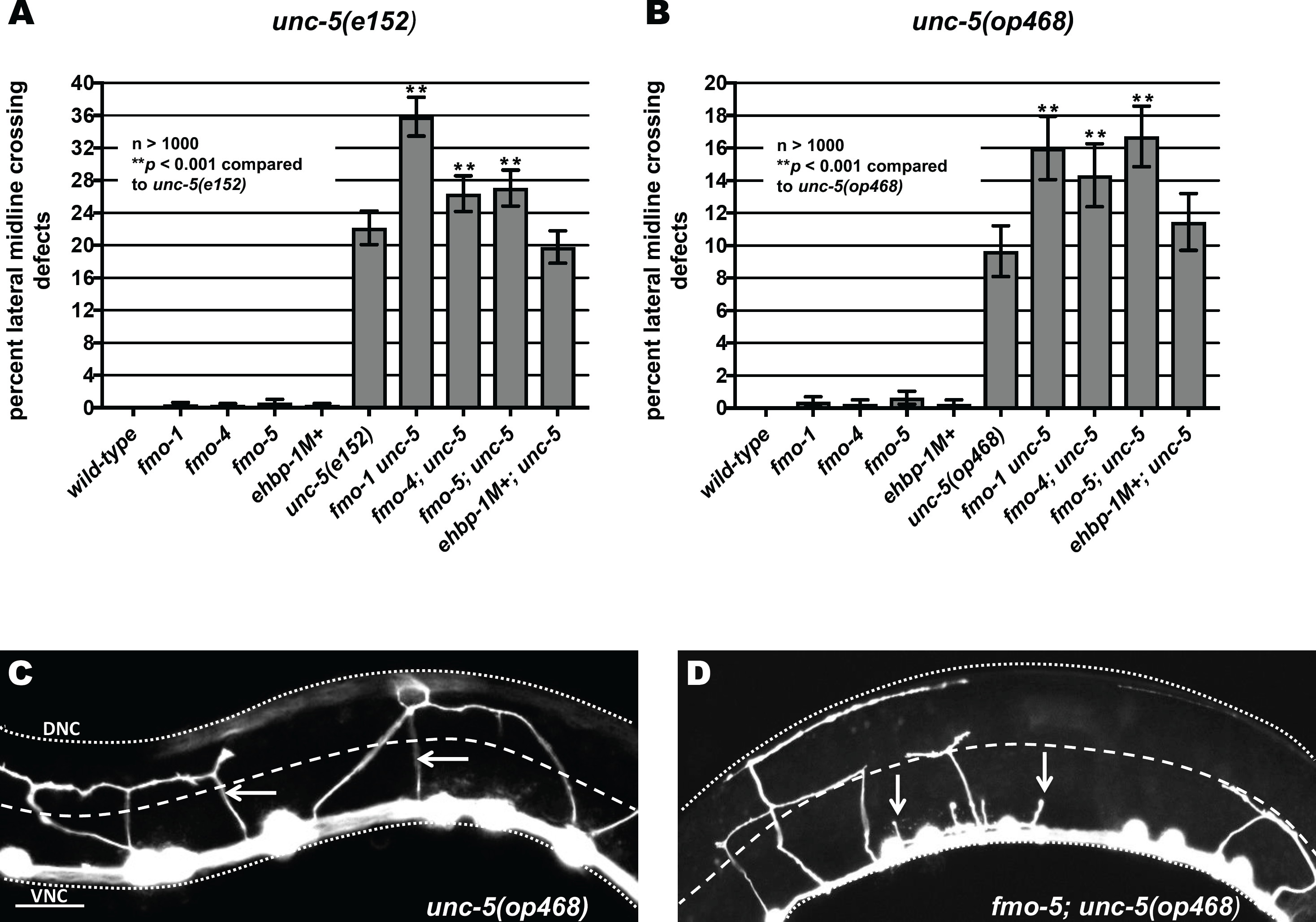
Axon pathfinding defects of hypomorphic *unc-5* mutants are enhanced by loss of *fmo-1* and *fmo-4*. **(A) and (B)** Quantification of VD/DD axons that failed to cross the lateral midline of L4 hermaphrodites in hypomorphic *unc-5(e152)* and *unc-5(op468)* mutants alone and in double mutant animals. Error bars represent 2x standard error of the proportion; double asterisks (**) indicates a significant difference between *unc-5(e152)* or *unc-5(op468)* alone and the double mutants (*p* < 0.001) determined by Fisher’s exact test. Only visible commissural processes emanating from the ventral nerve cord were scored. **(C,D)** Fluorescence micrographs of VD/DD axons (arrows) in L4 hermaphrodites. The lateral midline of the animal is indicated by the dashed white line. The dorsal nerve cord and ventral nerve cord are indicated by dotted white lines. Dorsal is up, anterior is to the left. Scale bar represents 5μm. (C) In the weak loss of function *unc-5(op468)* mutants, axons crossing the lateral midline are indicated (arrows). (D) In *fmo-5(tm2438); unc-5(op468)*, some axons cross the lateral midline, but many terminate before crossing the lateral midline (arrows).

### *fmo-1, fmo-4* and *fmo-5* act cell-autonomously in the VD/DD neurons

Expression of the *fmo-1, fmo-4* and *fmo-5* coding regions were driven in VD/DD motor neurons using the *unc-25* promoter. *Punc-25::fmo* transgenes significantly rescued lateral midline crossing defects in *fmo; unc-5(op468)* and *fmo; unc-5(e152)* (Figure 6). These data suggest that the axon defects observed in *fmo* mutants are due to mutation of the *fmo* genes themselves, and that *fmo-1, 4,* and *5* can act cell-autonomously in the VD/DD neurons in axon guidance.

**Fig. 6.**
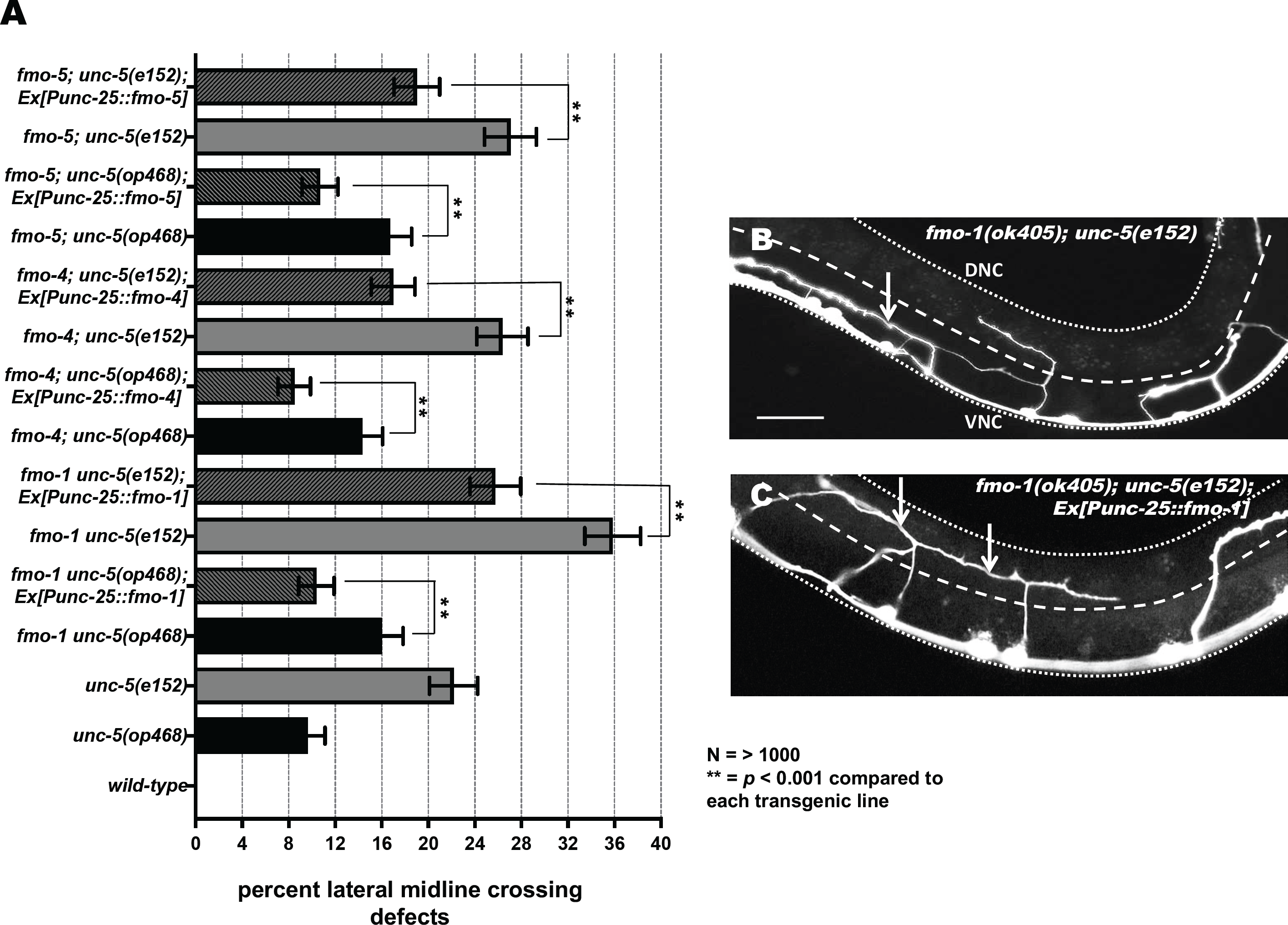
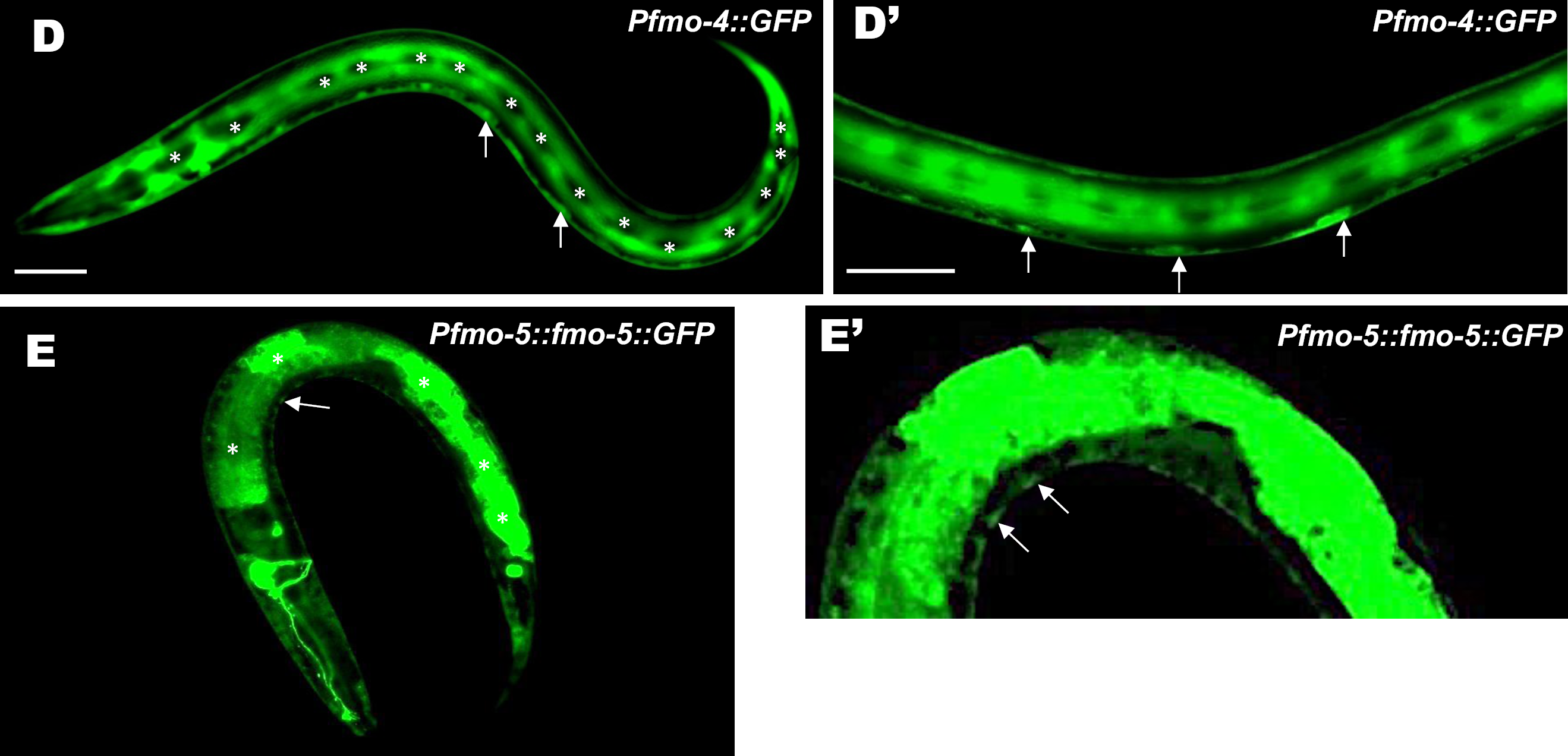
Expression of *fmo-1, fmo-4 and fmo-5* in VD/DD neurons rescues axon pathfinding defects. **(A)** The percentages of VD/DD axons failing to cross the lateral mid-line are as described in Figure 5A. *unc-5* double mutant genotypes are indicated, and the *Punc-25::fmo-1, Punc-25::fmo-4,* and *Punc-25::fmo-5* transgenes are bracketed. Data for transgenic strains are the combined results from three independently-derived transgenes with similar effects. Double asterisks (**) indicate a significant difference between the double mutant and the transgenic strain (*p* <0.001; Fisher’s exact test). Error bars represent 2x standard error of the proportion. **(B, C)** Micrographs of mutant and rescued animals. Dorsal is up, anterior to the left. Scale bar represents 5μm. The lateral midline of the animal is indicated by the dashed white line. The dorsal nerve cord and ventral nerve cord are indicated by dotted white lines. (B) *fmo-1(ok405) unc-5(e152)* axons often fail to cross the lateral midline (arrow). (C) *fmo-1(ok405) unc-5(e152*); Ex(*Punc-25::fmo-1*) axons crossed the lateral midline (arrows). **(D-E and D’-E’)** Images are micrographs of L2 animals with transgenic expression of *Pfmo-4::gfp* and *Pfmo-5::fmo-5::gfp*. Dorsal is up and anterior is left. Scale bar: 5μm. (D) *fmo-4::gfp* is broadly expressed, including in hypodermis and in cells along the ventral nerve cord that resemble motor neurons. Expression is not evident in the lateral hypodermal seam cells (asterisks). (D’) Enlarged image of *fmo-4::gfp* expression in ventral nerve cord cells (arrows). (E). *fmo-5::gfp* is expressed strongly in the gut (asterisks), as well as in cells along the ventral nerve cord. (arrow) (E’) Enlarged image of *fmo-5::gfp* expression in ventral nerve cord cells (arrows).

Previous studies showed that *fmo-1* and *fmo-5* promoter regions were active in intestinal cells and the excretory gland cell, whereas the *fmo-4* promoter was active in hypodermal cells, duct and pore cells [32, 36]. *ehbp-1* is expressed in all somatic cells including neurons [33]. Furthermore, cell-specific transcriptome profiling indicated that *fmo-1, fmo-4* and *fmo-5* were expressed in embryonic and adult neurons, including motor neurons [37-39]. We fused the upstream promoter regions of *fmo-1, fmo-4,* and *fmo-5* to *gfp*. We could observe no *fmo-1::gfp* expression in transgenic animals, in contrast to previous studies using a *LacZ* reporter [32]. However, transcriptome profiling indicates neuronal expression of *fmo-1 [39].* Our *fmo-1::gfp* transgene might be missing regulatory regions required for expression*. fmo-4::gfp* was expressed strongly in hypodermal cells, excluding the seam cells and vulval cells, consistent with previous studies [32] (Figure 6D). We also observed *fmo-4::gfp* expression in cells in the ventral nerve cord (Figure 6 D and D’). *Pfmo-5::fmo-5::gfp* was expressed strongly in the intestine as previously reported [32] (Figure 6E). We also observed expression along the ventral nerve cord (Figure 6E and E’).

In sum, previous expression studies combined with those described here suggest that *fmo-1,4,5* and *ehbp-1* are expressed in neurons, and that *fmo-1,4,* and *5* can act cell-autonomously in the VD/DD motor neurons in axon guidance.

### *fmo-1, fmo-4* and *fmo-5* mutants display increased growth cone filopodial protrusion

The growth cones of dorsally-directed VD commissural axons are apparent in early L2 larvae (Figure 2B). We imaged VD growth cones at 16 hours post-hatching, when the VD growth cones have begun their dorsal migrations, as described previously [15]. *fmo-1, fmo-4* and *fmo-5* mutant growth cones displayed longer filopodial protrusions compared to wild type (e.g. 0.96 μm in wild type compared with 1.55 μm in *fmo-5(tm2438)*; *p < 0.001*) (Figure 7). This effect was not significant in *ehbp-1(ok2140)* (Figure 7). Growth cone area was not significantly different in any mutant. These results suggest that *fmo-1, fmo-4* and *fmo-5* normally limit growth cone filopodial protrusion length. This is consistent with ectopic axon branches observed in post-development VD/DD neurons in these mutants (Figure 3), as other mutants with increased growth cone filopodial protrusions (e.g. *unc-5, unc-73, unc-33*) also display ectopic branches, likely due to failure of filopodial retraction and subsequent consolidation into a neurite [15, 17].

**Fig. 7.**
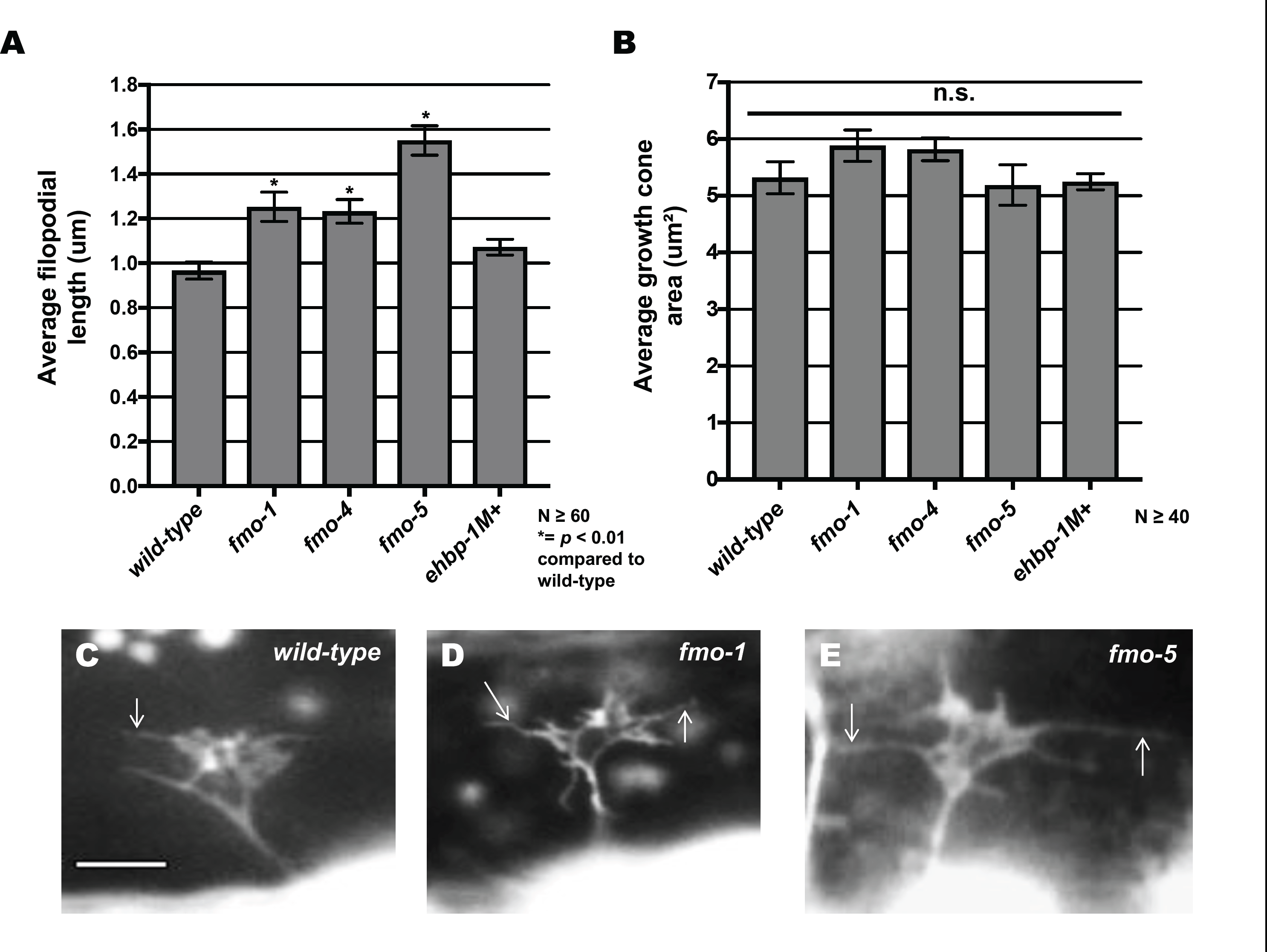
Mutations in *fmo-1, fmo-4* and *fmo-5* increase VD growth cone filopodial length. **(A,B)** Quantification of VD growth cone filopodial length and growth cone area in wild-type and mutant animals. (A) Average filopodial length, in μm. (B) Growth cone area in μm^2^. Error bars represent 2x standard error of the mean; asterisks indicate the significant difference between wild-type and the mutant phenotype (\**p* < 0.01) determined by two-sided *t*-test with unequal variance. n.s., not significant. **(C-E)** Fluorescence micrographs of VD growth cones; (C) A wild-type VD growth cone. (D) *fmo-1(ok405)* and (E) *fmo-5(tm2438) g*rowth cones showing increased filopodial protrusion in the form of longer filopodia. Arrows indicate representative filopodia. Scale bar: 5μm.

### *fmo-1, fmo-4, fmo-5* and *ehbp-1* mutations suppress activated *myr::unc-40* and *myr::unc-5* and activated Rac GTPases

Previous studies showed that UNC-6/netrin signaling via the heterodimeric UNC-40/UNC-5 receptor leads inhibition of growth cone protrusion important in UNC-6/Netrin’s role in repulsive axon guidance [15, 17]. Constitutive activation of this pathway using expression of myristoylated versions of the cytoplasmic domains of UNC-40 and UNC-5 (*myr::unc-40* and *myr::unc-5*) results in small growth cones with few if any filopodial protrusions (i.e. protrusion is constitutively inhibited by MYR::UNC-40 and MYR::UNC-5) [15, 17, 18]. Loss of *fmo-1, fmo-4, fmo-5* and *ehbp-1* significantly suppressed inhibition of filopodial protrusion and growth cone size caused by *myr::unc-40* (Figure 8) and *myr::unc-5* (Figure 9).

**Fig. 8.**
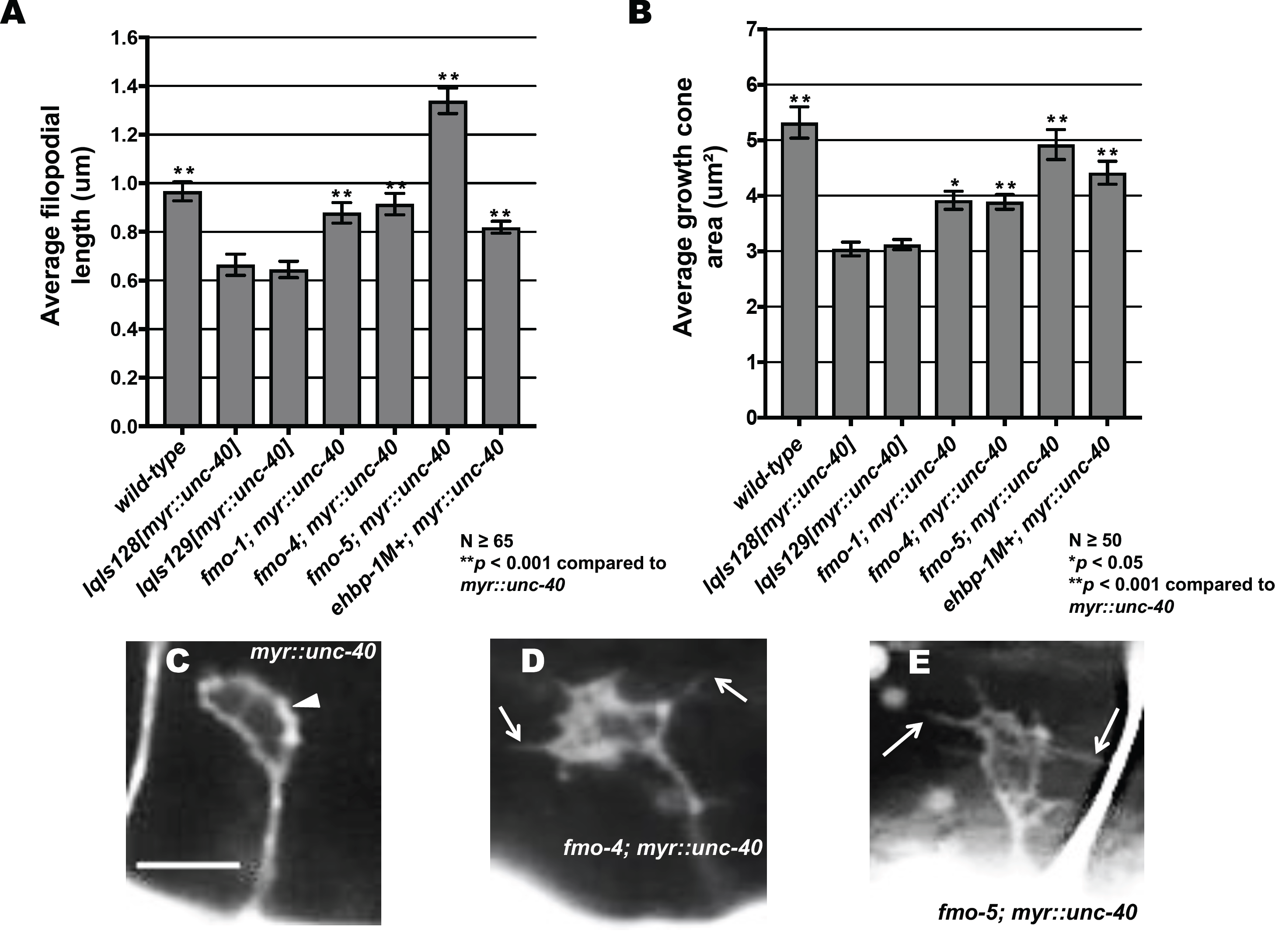
FMO-1, FMO-4, FMO-5, and EHBP-1 are required for MYR::UNC-40-mediated inhibition of VD growth cone protrusion. **(A,B)** Quantification of VD growth cone filopodial length and growth cone area in *wild-type, myr::unc-40 (lqIs128* and *lqIs129)* and double mutants. (A) Average filopodial length, in μm. **(B)** Growth cone area in μm^2^. Error bars represent 2x standard error of the mean. Asterisks indicate significant difference between *myr::unc-40, wild-type* and the double mutants (\**p* < 0.05, ** *p* < 0.001) determined by two-sided *t*-test with unequal variance. **(C-E)** Fluorescent micrographs of mutant VD growth cones; (C) Image of a *myr::unc-40* growth cone in an early L2 animal. The arrowhead points to a growth cone with little or no filopodial protrusion. (D, E) Images of *fmo-4(ok294); myr::unc-40* and *fmo-5(tm2438); myr::unc-40* growth cones. Filopodial protrusions are indicated (arrows). Scale bar: 5μm. *fmo-1(ok405); myr::unc-40* double mutants were built and compared with *lqIs129[myr::unc-40]* due to the linkage of the *lqIs128* transgene.

**Fig. 9.**
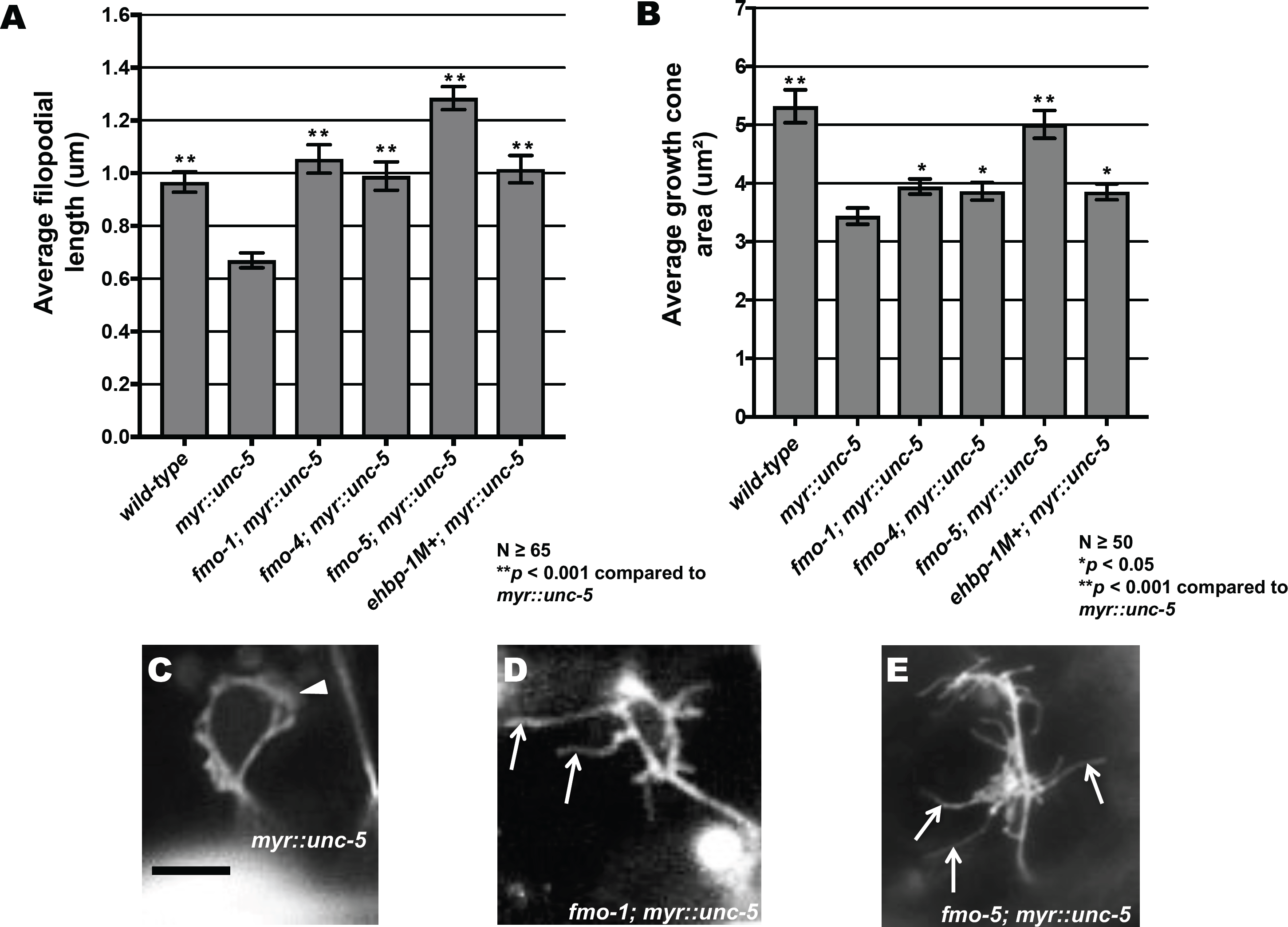
FMO-1, FMO-4, FMO-5, and EHBP-1 are required for MYR::UNC-5-mediated inhibition of VD growth cone protrusion. **(A,B)** Quantification of VD growth cone filopodial length and growth cone area in *wild-type, myr::unc-5,* and double mutants. (A) Average filopodial length, in μm. (B) Growth cone area in μm^2^. Error bars represent 2x standard error of the mean. Asterisks indicate significant difference between *myr::unc-5* and the double mutants (\**p* < 0.05, \*\**p* < 0.001) determined by two-sided *t*-test with unequal variance. **(C-E)** Representative fluorescent micrographs of mutant VD growth cones; (C) Image of a *myr::unc-5* growth cone in an early L2 animal. The arrowhead points to a growth cone with limited protrusion. (D, E) Images of *fmo-1(ok405); myr::unc-5* and *fmo-5(tm2438); myr::unc-5* growth cones. Arrows point to filopodial protrusions. Scale bar: 5μm.

Expression of activated CED-10(G12V) and MIG-2(G16V) in the VD neurons results in reduced growth cone protrusion similar to MYR::UNC-40 and MYR::UNC-5 [17]. We found that *fmo-1, fmo-4* and *fmo-5* suppressed filopodial protrusion deficits caused by *ced-10(G12V)* and *mig-2(G16V)* (Figure 10). *ehbp-1* suppressed *mig-2(G16V)*, but *ehbp-1(ok2140M+);ced-10(G12V)* double mutants were inviable and could not be scored. Furthermore, *fmo-4* and *fmo-5,* but not *fmo-1*, significantly suppressed growth cone size reduction caused by CED-10(G12V) and MIG-2(G16V). *ehbp-1* also suppressed growth cone size reduction of MIG-2(G16V). Taken together, these data indicate that functional FMO-1, FMO-4, FMO-5, and EHBP-1 are required for the full effect of MYR::UNC-40, MYR::UNC-5, CED-10(G12V), and MIG-2(G16V) on growth cone protrusion inhibition, including filopodial protrusion and growth cone size.

**Fig. 10.**
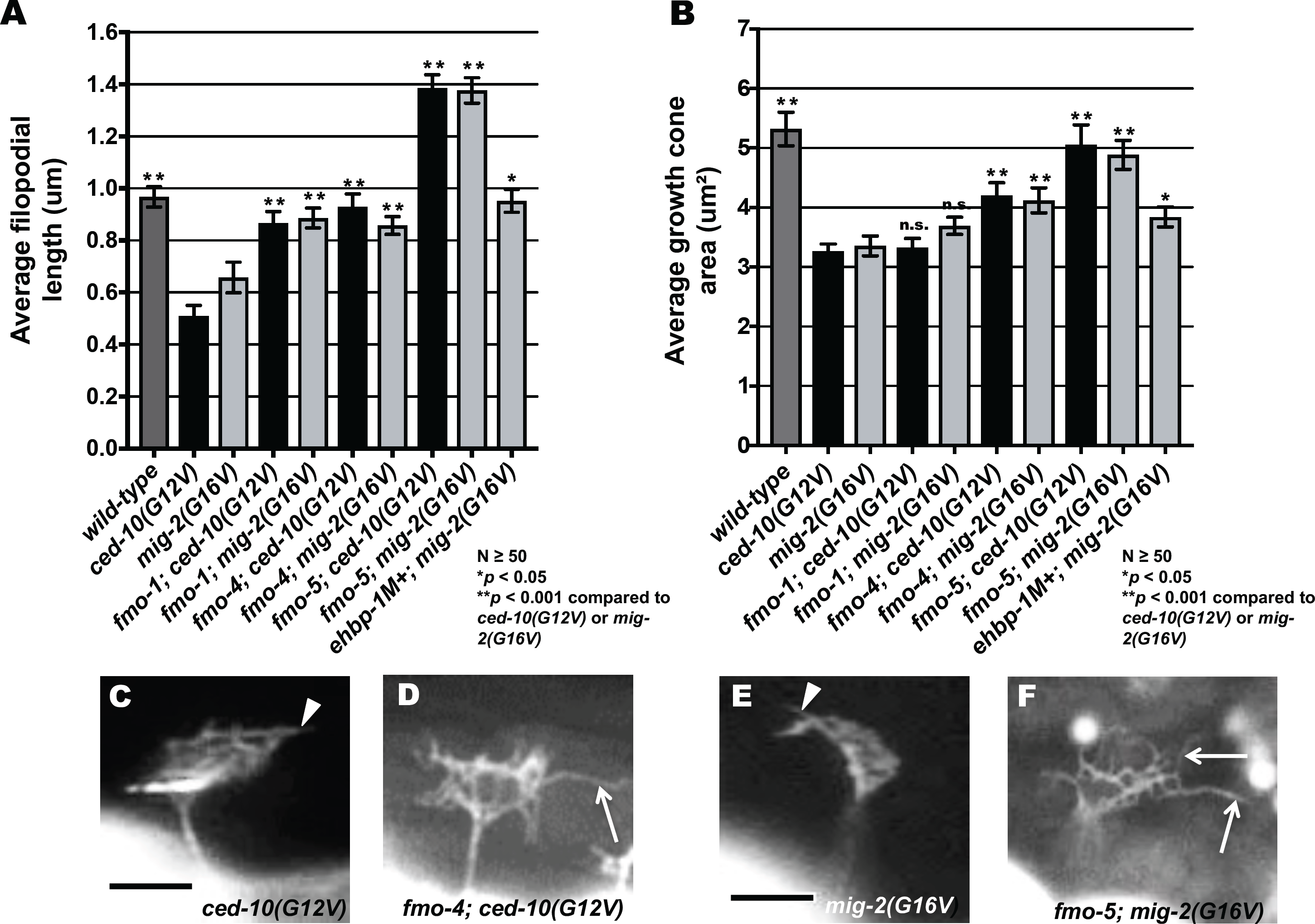
FMO-1, FMO-4, FMO-1, and EHBP-1 are required for Rac GTPase-mediated inhibition of VD growth cone protrusion. **(A,B)** Quantification of VD growth cone filopodial length and growth cone area in *wild-type*, activated *ced-12(G12V)* and *mig-2(G16V),* and double mutants. (A) Average filopodial length, in μm. (B) Growth cone area in μm^2^. Error bars represent 2x standard error of the mean. Asterisks indicate significant difference between *ced-10(G12V) mig-2(G16V)* and their respective double mutants (\**p* < 0.05, \*\**p* < 0.001) determined by two-sided *t*-test with unequal variance. n.s., not significant. **(C-F)** Representative fluorescent micrographs of mutant VD growth cones. (C,D) Images of *ced-10(G12V)* and *fmo-4(ok294); ced-10(G12V)* growth cones. The arrowhead in (C) points to a growth cone with limited protrusion, and the arrow in (D) indicates a filopodial protrusion. (E,F) Images of *mig-2(G16V)* and *fmo-5(tm2438); mig-2(G16V)* growth cones. The arrowhead in (E) points to a growth cone with limited protrusion, and the arrow in (F) indicates a filopodial protrusion. Scale bar: 5μm.

### FMO-5 can inhibit growth cone protrusion

*fmo-5* loss-of-function mutant growth cones displayed excessively protrusive filopodia (Figure 7) and suppressed activated UNC-40/UNC-5 and Rac signaling (Figures 8-10). Transgenic expression of wild-type FMO-5 driven by its endogenous promoter rescued the long filopodial protrusions seen in *fmo-5(tm2438)* mutant VD growth cones (Figure 11A-D). In a wild-type background, *fmo-5* transgenic expression resulted in growth cones with smaller area and shortened filopodia (Figure 11E, F and H), indicating that wild-type FMO-5 activity can inhibit growth cone protrusion. This was not observed in the *fmo-5(tm2438)* background, possibly due to the decreased levels of FMO-5 compared to the wild-type background.

**Fig. 11.**
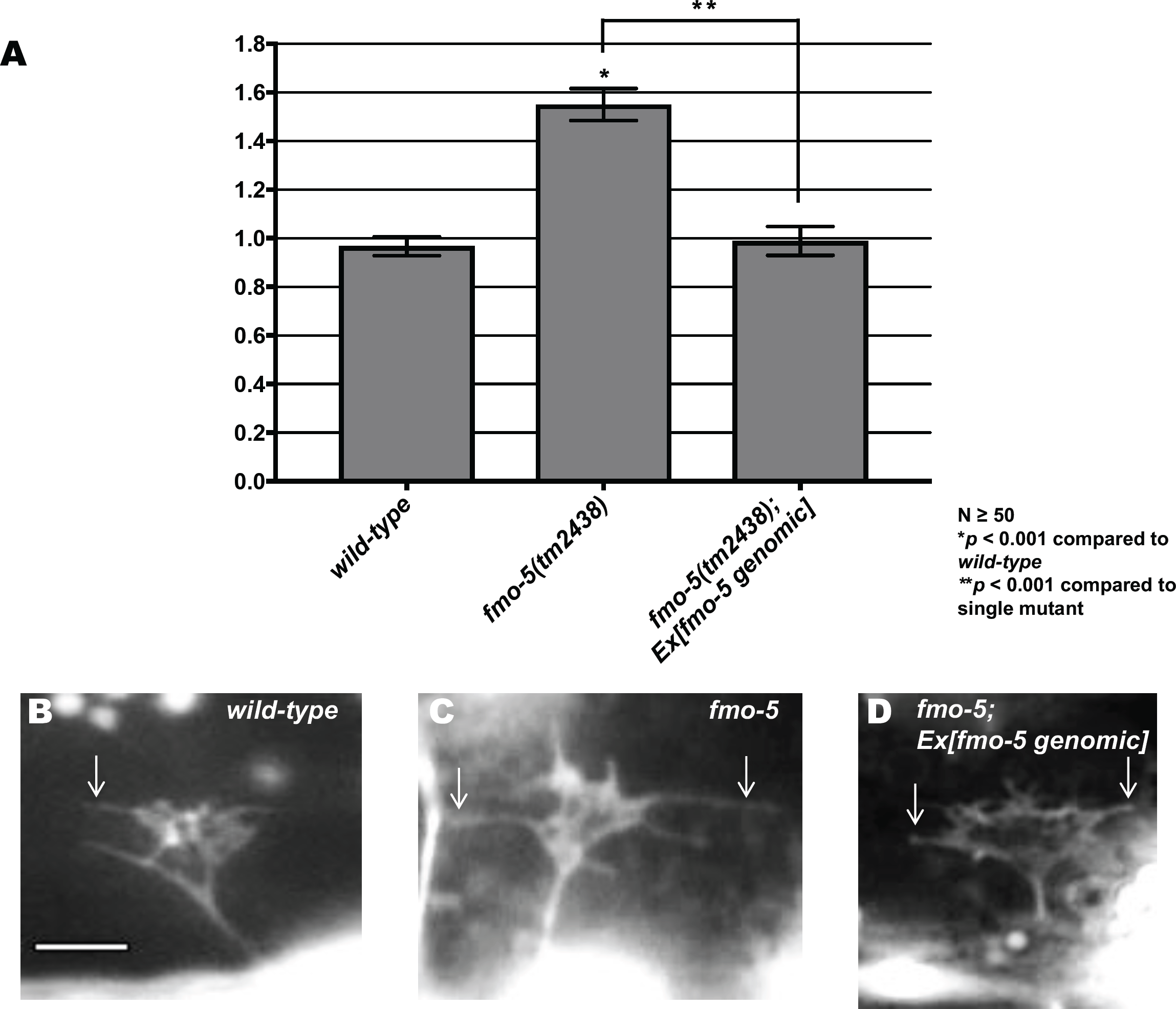
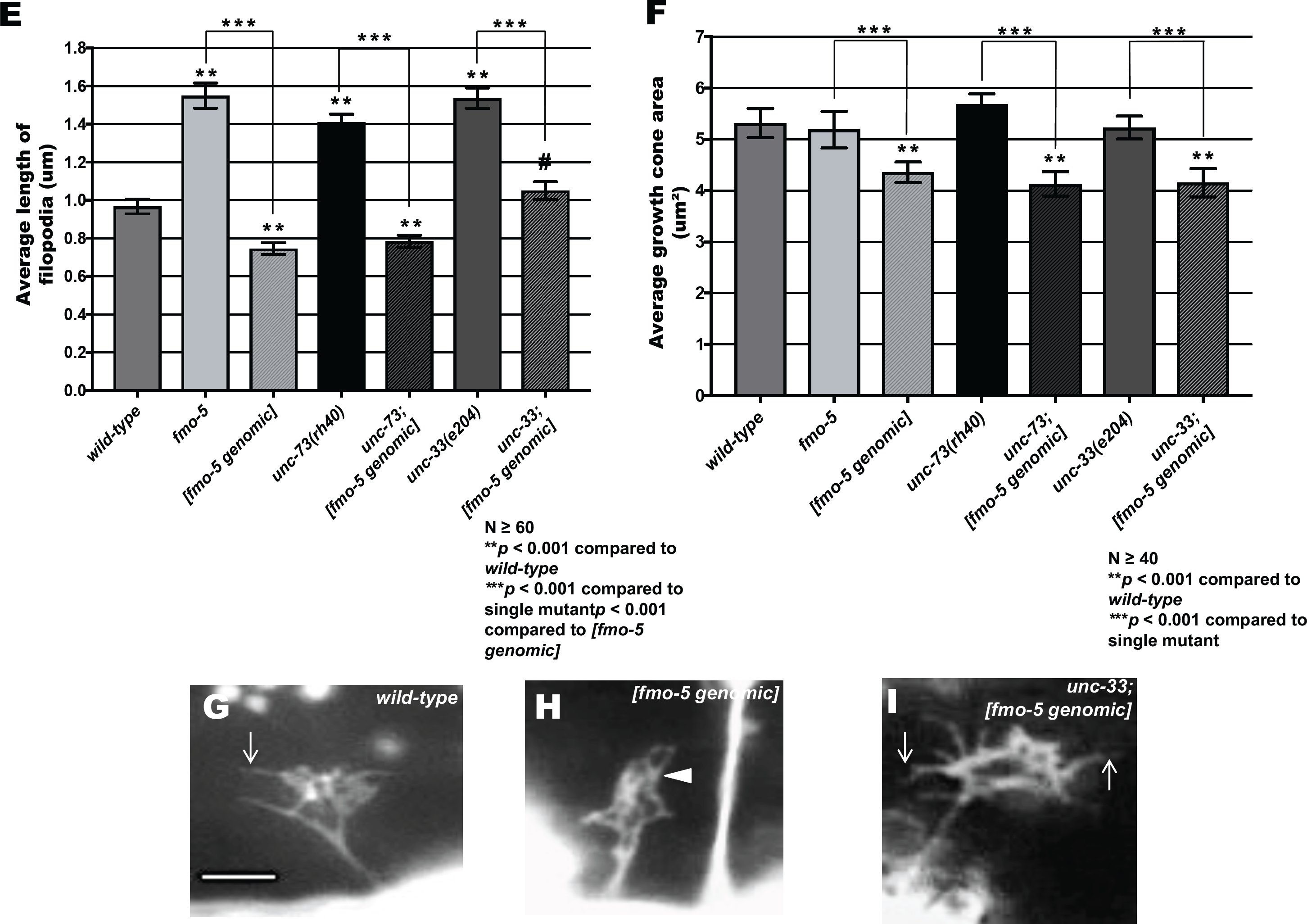
FMO-5 can inhibit growth cone filopodial protrusion. **(A**) Rescue of *fmo-5(tm2438)* VD growth cone filopodial protrusions by transgenes containing genomic *fmo-5* (*Ex[fmo-5 genomic]*). Data for transgenic arrays are the combined results from three independently-derived arrays with similar effects. Average lengths of filopodial protrusions are shown (μm). Error bars represent 2x standard error of the mean. Single asterisks (*) indicate a significant difference between wild type and the mutant (*p* < 0.001); Double asterisks (**) indicates a significant difference between the mutant and rescuing transgene (*p* < 0.001) determined by two-sided *t*-test with unequal variance **(B-D)** Fluorescence micrographs of VD growth cones in *wild-type, fmo-5(tm2438)*, and *fmo-5(tm2438); Ex[fmo-5 genomic]*. Arrows indicate representative filopodia. Scale bar: 5μm. **(E,F)** Quantification of VD growth cone filopodial length and growth cone area in indicated genotypes. Error bars represent 2x standard error of the mean. Asterisks indicate significant difference between wild-type and mutants (** *p* < 0.001) and *** indicate a significant difference between each single mutant compared to the double mutant. Pound signs (**#)** indicate a significant difference between [*fmo-5 genomic*] and double mutant (#*p* < 0.001) determined by two-sided *t*-test with unequal variance. **(G-I)** Fluorescence micrographs of VD growth cones from *wild-type, [fmo-5 genomic]*, and *unc-33; [fmo-5 genomic]*. The arrowhead points to a growth cone with limited protrusion. Arrows indicate representative filopodia. Scale bar: 5μm.

**Fig. 12.**
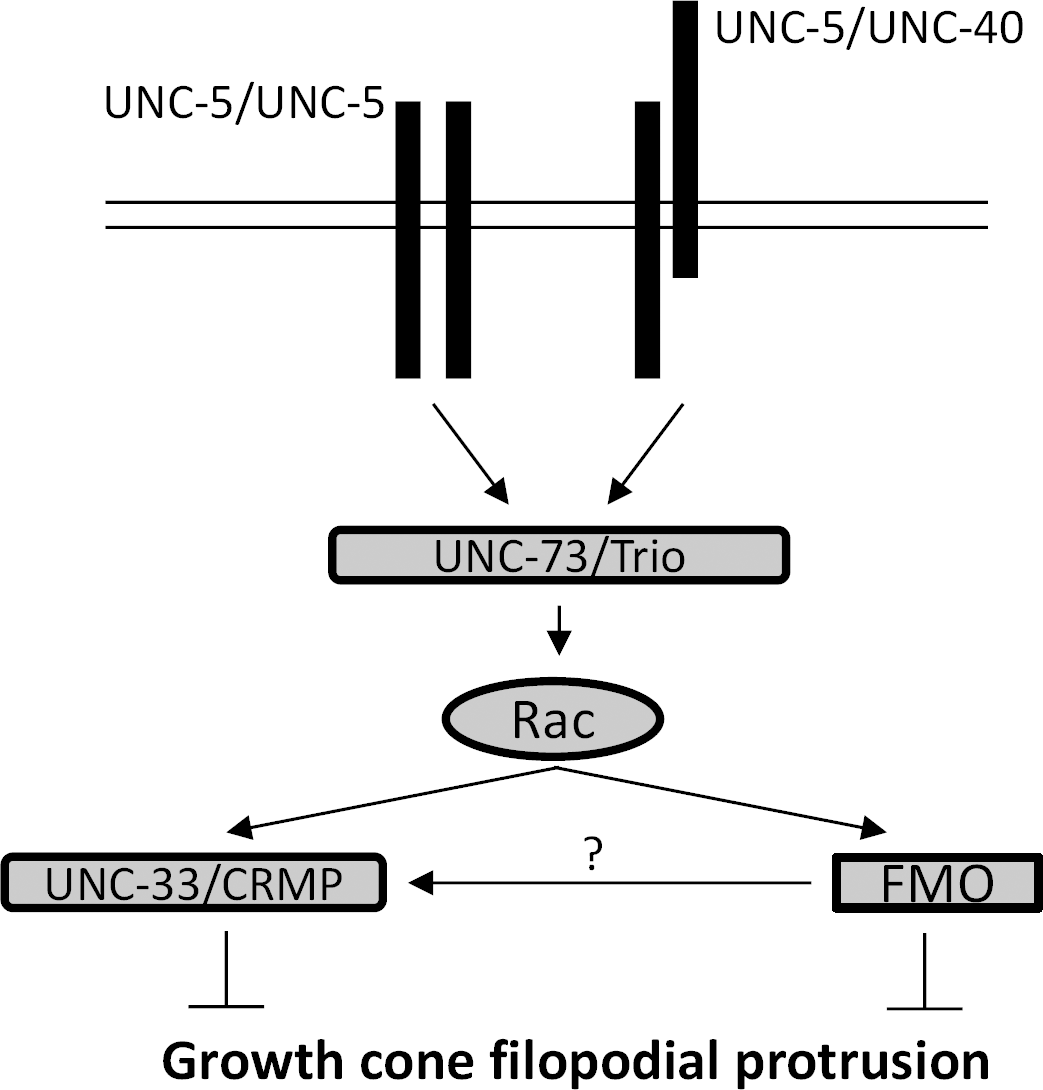
Genetic model of inhibition of growth cone protrusion. UNC-5 homodimers and/or UNC-40/UNC-5 heterodimers act through the Rac GTP exchange factor UNC-73/Trio and the Rac GTPases, which then utilize the flavin monooxygenases and UNC-33/CRMP to inhibit protrusion. The FMOs might inhibit protrusion directly, by possibly directly oxidizing F-actin, or by promoting phosphorylation of UNC-33/CRMP.

A pathway involving the Rac GTPases MIG-2 and CED-10, the Rac GEF UNC-73/Trio, and the putative cytoskeletal interacting molecule UNC-33/CRMP act downstream of MYR::UNC-40 to inhibit protrusion [17]. In this pathway, UNC-33/CRMP acts downstream of the Rac GTPases [17], similar to the FMOs and EHBP-1 described in Figure 10. *unc-73(rh40)* specifically attenuates the Rac GEF domain of UNC-73 and results in excessively-protrusive growth cones, including increased growth cone area and filopodial length [17] (Figure 11F). Transgenic *fmo-5* expression in *unc-73(rh40)* resulted in inhibited growth cone area and filopodial length compared to *unc-73(rh40)* alone, suggesting that FMO-5 can inhibit protrusion in the absence of UNC-73 Rac GEF activity. Transgenic *fmo-5* expression inhibited growth cone size in *unc-33(e204)*, but had a reduced capacity to inhibit filopodial protrusion (i.e. filopodial protrusion in *unc-33(e204)* with transgenic *fmo-5* expression was reduced compared to *unc-33* alone but was increased relative to *fmo-5* transgenic expression alone) (Figure 11E-I). In sum, these data indicate that that FMO-5 (and possibly FMO-1 and FMO-4) act downstream of the Rac GTPases MIG-2 and CED-10 in filopodial inhibition. FMO-5 might also act downstream of UNC-33, but the hybrid interaction of *fmo-5* transgenic expression with *unc-33(e204)* mutants suggests that FMO-5 and UNC-33 might represent distinct pathways downstream of the Rac GTPases to inhibit filopodial protrusion.

## Discussion

Results here implicate the *C. elegans* flavoprotein monooxygenase molecules FMO-1, FMO-4 and FMO-5 in inhibition of growth cone protrusion via UNC-6/Netrin receptor signaling in repulsive axon guidance. The MICAL molecule found in vertebrates and *Drosophila* is a flavoprotein monooxygenase required for semaphorin-plexin mediated repulsive motor axon guidance [27, 40]. MICAL is a multi-domain molecule that also includes a calponin homology (CH) domain, a LIM domain and multiple CC domains. No molecule encoded in the *C. elegans* genome has a similar multi-domain organization. However, the Eps-15 homology (EH) domain binding protein EHBP-1 is similar to the non-FMO portion of MICAL and contains a CH domain [33]. We show here that EHBP-1 also is also involved in inhibition of growth cone protrusion and axon guidance. Thus, while *C. elegans* does not have a multidomain MICAL-like molecule, it is possible that the functional equivalents are the FMOs and EHBP-1.

**FMO-1, FMO-4, FMO-5 and EHBP-1 regulate axon guidance and growth cone filopodial protrusion.** *fmo-1, fmo-4, fmo-5,* and *ehbp-1* mutants display defects in dorsal guidance of the VD/DD motor axons that are repelled from UNC-6/Netrin (Figure 3). Double mutant analysis did not uncover significant redundancy, suggesting that these molecules might have discrete roles in axon guidance. Consistent with this idea, *fmo-4* and *ehbp-1* mutually suppress VD/DD axon guidance defects. *fmo-2* and *fmo-3* mutations displayed no significant defects alone, suggesting that they are not involved in axon guidance. *fmo-2* did significantly enhance *fmo-4*. Possibly, *fmo-2* and *fmo-3* have roles in axon guidance that were not revealed by the mutations used.

*Drosophila* and vertebrate MICAL regulate actin cytoskeletal dynamics in both neuronal and non-neuronal processes through direct redox activity of the monooxygenase domain [27, 30, 41-45]. In *Drosophila*, loss of MICAL showed abnormally shaped bristles with disorganized and larger F-actin bundles, whereas, overexpression of MICAL caused a rearrangement of F-actin into a complex meshwork of short actin filaments [29]. Here we show that loss of *fmo-1, fmo-4,* and *fmo-5* resulted in longer filopodial protrusions in the VD motor neurons (Figure 7), suggesting that their normal role is to limit growth cone filopodial protrusion. Indeed, transgenic expression of wild-type FMO-5 resulted in VD growth cones with a marked decrease in growth cone filopodial protrusion (Figure 11). Growth cone size was not affected in any loss-of-function mutation, but growth cone size was reduced by transgenic expression of wild-type FMO-5 (Figure 11), suggesting a role of the FMO-5 in both filopodial protrusion and growth cone lamellipodial protrusion.

Previous studies have shown that *Drosophila* MICAL may require both its FMO and CH domain to induce cell morphological changes; however, mammalian MICAL in non-neuronal cell lines requires only its FAD domain suggesting a difference in the mechanism of action in these MICALs [29, 46]. These data suggest that in some cases, the FMO domain is sufficient for the function of MICAL. Thus, single domain FMOs as in *C. elegans* could function despite lacking the multi-domain structure of MICAL. Loss of EHBP-1, which contains a CH domain and is similar to the non-FMO portion of MICAL (Figure 1), also resulted in VD/DD axon guidance defects, but did not significantly affect growth cone filopodial protrusion. EHBP-1 might act with the FMOs in axon guidance. Phenotypic differences could be due to EHBP-1-dependent and independent events, or to the wild-type maternal contribution in *ehbp-1* homozygous mutants derived from a heterozygous mother. It is also possible that EHBP-1 affects axon guidance independently of the FMOs. EHBP-1 is involved in Rab-dependent endosomal vesicle trafficking by bridging interaction of endosomal Rabs with the actin cytoskeleton [33, 47]. MICAL has also been implicated in Rab-dependent endosomal biogenesis and trafficking [48-50], suggesting that FMO/EHBP-1 and MICALs might share common functions, although it remains to be determined if FMOs in *C. elegans* regulate endosomal trafficking.

MICAL has been shown to directly oxidize cysteine residues in F-actin, leading to actin depolymerization and growth cone collapse [29, 30, 51, 52]. We speculate that FMO-1, FMO-4, and FMO-5 might act by a similar mechanism to inhibit growth cone filopodial protrusion. The role of EHBP-1 is less clear, but previous studies have shown that *Drosophila* MICAL might require both its FMO and CH domain to induce cell morphological changes [29]. Thus, in axon guidance, FMO-1, FMO-4, and FMO-5 might require the CH domain provided by EHBP-1 in some instances. Mammalian MICAL requires only the FMO domain [46], suggesting that in some cases the CH domain is not required and the FMO domain can act alone. Future studies will be directed at answering these questions.

### FMOs can act autonomously in the VD/DD neurons

Expression of full length *fmo-1, fmo-4* and *fmo-5* coding regions under the control of the *unc-25* promoter specific for GABA-ergic neuron expression (including the VD/DD neurons) rescued VD/DD axon guidance defects (Figure 6). Furthermore, the promoters of *fmo-4* and *fmo-5* were active in ventral nerve cord cells (Figure 6). Cell-specific transcriptome profiling indicated that *fmo-1, fmo-4* and *fmo-5* were expressed in embryonic and adult neurons, including motor neurons [37-39]. Together, these results suggest that the FMOs can act cell-autonomously in the VD/DD neurons in axon guidance.

### FMO-1, FMO-4 and FMO-5 mediate UNC-6/Netrin receptor signaling in growth cone inhibition of protrusion

Our findings suggest that the FMOs act with the UNC-40 and UNC-5 receptors to mediate UNC-6/netrin repulsive axon guidance and inhibition of growth cone protrusion. *fmo-1, fmo-4,* and *fmo-5* mutations enhanced axon pathfinding defects in *unc-40* and hypomorphic *unc-5* mutants (Figures 4 and 5). *ehbp-1* did not enhance *unc-40* or *unc-5*, suggesting discrete roles of these molecules or wild-type maternal *ehbp-1* contribution. However, *fmo-1, fmo-4, fmo-5*, and *ehbp-1* mutations each suppressed the effects of activated MYR::UNC-40 and MYR::UNC-5 on inhibition of growth cone protrusion (Figure 9). In this case, both filopodial protrusion and growth cone area was restored, consistent with a role of these molecules in inhibiting both growth cone filopodial and lamellipodial protrusion. That the FMOs and EHBP-1 were required for the effects of the constitutively active MYR::UNC-40 and MYR::UNC-5 suggest that they act downstream of these molecules in growth cone inhibition of protrusion.

### FMOs and EHBP-1 act downstream of Rac GTPase signaling in inhibition of growth cone protrusion

Similar to activated MYR::UNC-40 and MYR::UNC-5, constitutively-activated Rac GTPases CED-10(G12V) and MIG-2(G16V) inhibit VD growth cone protrusion. We show that *fmo-1, fmo-4, fmo-5* and *ehbp-1* mutations suppressed activated CED-10(G12V) and MIG-2(G16V) (e.g. double mutant growth cones displayed longer filopodial protrusions similar to *fmo-1, fmo-4, fmo-5* and *ehbp-1* single mutants) (Figure 10). Furthermore, loss of the Rac GTP exchange factor UNC-73/Trio had no effect on the inhibited growth cone phenotype of FMO-5 transgenic expression (i.e. the growth cones resembled those of *fmo-5* over expression alone) (Figure 11). UNC-73/Trio acts with the Rac GTPases CED-10 and MIG-2 in growth cone protrusion inhibition, and *unc-73* mutants display excessive growth cone protrusion [17]. That FMO-5 transgenic expression could inhibit protrusion in the absence of the Rac activator UNC-73/Trio suggests that FMO-5 acts downstream of UNC-73/Trio, consistent with the FMOs and EHBP-1 acting downstream of the Rac GTPases.

### FMO-5 might act upstream of UNC-33/CRMP

Previous studies have shown that the *C. elegans* CRMP-like molecule UNC-33 is required in a pathway downstream of Rac GTPases for inhibition of growth cone protrusion in response to UNC-6/Netrin [17]. *unc-33* loss-of-function mutants with FMO-5 transgenic expression displayed a mutually-suppressed phenotype. The excessively-long filopodial protrusions of *unc-33* mutants were reduced to wild-type levels, but were significantly longer than in animals with FMO-5 transgenic expression, and the growth cone area was reduced to resemble FMO-5 transgenic expression alone (Figure 11). This phenotype could be interpreted as FMO-5 acting upstream of UNC-33/CRMP (i.e. UNC-33/CRMP is required for the full effect on FMO-5 overexpression). Alternatively, this hybrid phenotype could be interpreted as FMO-5 and UNC-33/CRMP acting independently to inhibit protrusion.

One proposed mechanism of cytoskeletal regulation by MICAL is the production of the reactive oxygen species (ROS) H_2_O_2_ by the FAD domain in the presence of NADPH [53]. Upon activation by Sema3A, MICALs generate H_2_O_2_, which can, via thioredoxin, promote phosphorylation of CRMP2 by glycogen synthase kinase-3, leading to microtubule growth cone collapse [54]. This is consistent with our genetic results suggesting that FMO-5 may function upstream of UNC-33/CRMP in modulating the cytoskeleton of the VD growth cones to inhibit growth cone filopodial protrusion. CRMPs have been shown coordinate both microtubules and actin in axon elongation and growth cone dynamics [55, 56]. Thus, the FMOs have the potential to inhibit growth cone protrusion through direct oxidation of F-actin resulting in depolymerization, and through redox regulation of the activity of UNC-33/CRMP.

### Conclusion

In summary, we present evidence of a novel role of the *C. elegans* flavin-containing monooxygenase molecules (FMOs) in inhibition of growth cone protrusion downstream of UNC-6/Netrin signaling. The FMOs acted downstream of the UNC-6/Netrin receptors UNC-5 and UNC-40, and downstream of the Rac GTPases CED-10 and MIG-2. Future studies will determine if the FMOs regulate UNC-33/CRMP, if they cause actin depolymerization, or both, to inhibit growth cone protrusion.

## Materials and methods

### Genetic methods

Experiments were performed at 20**°**C using standard *C. elegans* techniques [57]. Mutations used were LGI: *unc-40(n324), unc-73(rh40)*; LGII: *juIs76[Punc-25::gfp]*; LGIII: *fmo-3(gk184651)*; LGIV: *fmo-1(ok405), fmo-2(ok2147), lqIs128[Punc-25::myr::unc-40], unc-5(op468* and *e152), unc-33(e204*); LGV: *fmo-4(ok294), fmo-5(tm2438), ehbp-1(ok2140M+)*; LGX: *lqIs182[Punc-25::mig-2(G16V)]*. Chromosomal locations not determined: *lqIs129[Punc-25::myr::unc-40], lqIs296[Punc-25::myr::unc-5], lqIs204[Punc-25::ced-10(G12V)], lhIs6[Punc-25::mCherry], lqIs311[fmo-5 genomic]* by integration of *lqEx1047*. Extrachromosomal arrays were generated using standard gonadal injection [58] and include: *lqEx901* and *lqEx931[Pehbp-1::gfp, Pgcy-32::yfp]; lqEx1014, lqEx1015, lqEx1016, lqEx1045, lqEx1046 and lqEx1047[Pfmo-5::fmo-5, Pgcy-32::yfp]; lqEx949, lqEx950, lqEx951, lqEx1053, lqEx1054* and *lqEx1055[Punc-25::fmo-1, Pgcy-32::yfp]; lqEx1057, lqEx1058* and *lqEx1060[Punc-25::fmo-4, Pgcy-32::yfp]; lqEx952, lqEx953, lqEx954, lqEx1061, lqEx1062, lqEx1063, lqEx1078, lqEx1079* and *lqEx1080[Punc-25::fmo-5, Pgcy-32::yfp]; lqEx1113* and *lqEx1114[Pfmo-5::fmo-5::GFP, Pstr-1::gfp]; whEx28[Pfmo-4::gfp, pRF4/rol-6]*. Multiple (≥3) extrachromosomal transgenic lines of *Pfmo-5::fmo-5* for overexpression data of *fmo-5* were analyzed with similar effect, and one was chosen for integration and further analysis. Genotypes containing M+ indicate that homozygous animals from a heterozygous mother were scored. The *ehbp-1(ok2140M+)* strain was balanced with the *nT1* balancer.

### Transgene construction

Details about transgene construction are available by request. *Punc-25::fmo-1, Punc-25::fmo-4* and *Punc-25::fmo-5* were made using the entire genomic regions of *fmo-1, fmo-4* and *fmo-5* respectively. Expression analysis for *fmo-5* was done by amplifying the entire genomic region of *fmo-5* along with its endogenous promoter (1.2kb upstream) and fusing it to *gfp* followed by the 3’ UTR of *fmo-5*.

### Analysis of axon guidance defects

VD neurons were visualized with a *Punc-25::gfp* transgene, *juIs76* [59], which is expressed in GABAergic neurons including the six DDs and 13 VDs, 18 of which extend commissures on the right side of the animal. The commissure on the left side (VD1) was not scored. In *wild-type*, an average of 16 of these 18 VD/DD commissures are apparent on the right side, due to fasciculation of some of the commissural processes (Figure 2C). In some mutant backgrounds, fewer than 16 commissures were observed (e.g. *unc-5*). In these cases, only observable axons emanating from the ventral nerve cord were scored for axon guidance defects. VD/DD axon defects scored include axon guidance (termination before reaching the dorsal nerve cord or wandering at an angle greater than 45° before reaching the dorsal nerve cord), lateral midline crossing (axons that fail to extend dorsally past the lateral midline) and ectopic branching (ectopic neurite branches present on the commissural processes). Fisher's exact test was used to determine statistical significance between proportions of defective axons. In double mutant comparisons, the predicted additive effect of the mutants was calculated by the formula P1+P2-(P1*P2), where P1and P2 are the phenotypic proportions of the single mutants. The predicted additive effect of single mutants was used in statistical comparison to the observed double mutant effect.

### Growth cone imaging

VD growth cones were imaged as previously described [15, 22]. Briefly, animals at 16 h post-hatching at 20°C were placed on a 2% agarose pad and paralyzed with 5mM sodium azide in M9 buffer, which was allowed to evaporate for 4 min before placing a coverslip over the sample. Some genotypes were slower to develop than others, so the 16 h time point was adjusted for each genotype. Growth cones were imaged with a Qimaging Rolera mGi camera on a Leica DM5500 microscope. Projections less than 0.5 μm in width emanating from the growth cone were scored as filopodia. Filopodia length and growth cone area were measured using ImageJ software. Filopodia length was determined by drawing a line from the base where the filopodium originates on the edge of the peripheral membrane to the tip of the filopodium. Growth cone area was determined by tracing the periphery of the growth cone, not including filopodial projections. Significance of difference was determined a two-sided *t-*test with unequal variance.

## Acknowledgments

The authors thank members of the Lundquist and Ackley labs for discussion, E. Struckhoff for technical assistance, and C. Dolphin for *fmo* strains and reagents. Some *C. elegans* strains were provided by the CGC, which is funded by NIH Office of Research Infrastructure Programs (P40OD010440). This work was supported by NIH grants R01NS040945, R56NS095682, and R21NS070417 to E.A.L., the University of Kansas Center for Molecular Analysis of Disease Pathways (NIH P20GM103638), and the Kansas Infrastructure Network of Biomedical Research Excellence (NIH P20GM103418), through which A.M.S. is a KINBRE Undergraduate Research Scholar.

